# Genes used together are more likely to be fused together in evolution by mutational mechanisms: A bioinformatic test of the used-fused hypothesis

**DOI:** 10.1101/2021.07.31.454590

**Authors:** Evgeni Bolotin, Daniel Melamed, Adi Livnat

## Abstract

Cases of parallel or recurrent gene fusions, whether in evolution or in cancer and genetic disease, are difficult to explain, as they require multiple of the same or similar breakpoints to repeat. The used-together-fused-together hypothesis holds that genes that are used together repeatedly and persistently in a certain context are more likely than otherwise to undergo a fusion mutation in the course of evolution—reminiscent of the Hebbian learning rule where neurons that fire together wire together. This mutational hypothesis offers to explain both evolutionary parallelism and recurrence in disease of gene fusions under one umbrella. Here, we test this hypothesis using bioinformatic data. Various measures of gene interaction, including co-expression, co-localization, same-TAD presence and semantic similarity of GO terms show that human genes whose homologs are fused in one or more other organisms are significantly more likely to interact together than random genes, controlling for genomic distance between genes. In addition, we find a statistically significant overlap between pairs of genes that fused in the course of evolution in non-human species and pairs that undergo fusion in human cancers. These results provide support for the used-together-fused-together hypothesis over several alternative hypotheses, including that all gene pairs can fuse by random mutation, but among pairs that have thus fused, those that have interacted previously are more likely to be favored by selection. Multiple consequences are discussed, including the relevance of mutational mechanisms to exon shuffling, to the distribution of fitness effects of mutation and to parallelism.

## Introduction

TRIM5 is a restriction factor that recognizes and inactivates retroviral capsids (*1*). CypA is a highly abundant cytosolic protein that, among other roles, potently binds several retroviral capsids, including HIV-1 (*2, 3*). Their respective genes fused at least twice independently by retroposition in two different simian lineages (*1, 4–9*), producing two similar new fused genes that provide resistance to certain lentiviruses (*4,5*). This is surprising: when observing a parallel point mutation, it is commonly assumed that random mutation hit the same base position twice. In contrast, in the case of a gene fusion such as described, multiple similar breakpoints defining the same two loci have to be drawn by chance twice. Mathematically, the latter is much harder to explain: if the probability of the former is low, the probability of the latter is negligible. Moreover, multiple other *TRIM* genes exist, a fusion of which to *CypA* would have likely also provided some, though smaller, resistance to retroviruses, as shown by *in vitro* studies (*10–12*); yet in both cases, *CypA* fused to *TRIM5* specifically (*1*).

According to a recent hypothesis, genes that are used together repeatedly and persistently in a certain context are more likely than otherwise to undergo a fusion mutation in the course of evolution (*13, 14*). In other words, “genes that are used together are fused together” (*14*). According to this hypothesis, genes that work together are likely transcribed at the same time and in the same place in the nucleus—for example in transcription factories, where DNA loops bring also distant collaborating genes together (*15–17*). This causes the chromatin to be open at both loci simultaneously, brings them close together spatially, and allows various downstream mechanisms, such as reverse transcription of the RNA, perhaps aided by trans-splicing, or other mechanisms (e.g., transposable-element mediated translocation, recombination, etc.) to generate a gene fusion (*13, 14*).

Although it has been known that genes interacting in one species are often fused in others (*18, 19*), and though it has been suggested that interaction can precede fusion in the course of evolution (*20, 21*), prior to ref. 14, the fusion of genes that work together in evolution had not been tied systematically to mutational mechanisms. One hypothesis had suggested that the (presumably random) fusion of two protein domains increases their effective concentrations with respect to each other and thus allows interactions between them to evolve (*18*). However, it was criticized by Doolittle, who questioned whether such a benefit is really provided by fusion, as genes do not need to be fused for their products to meet (*22*). In another realm— that of cancer research—it was argued that a pair of interacting genes expressed in the same transcription factory can undergo fusion at the RNA level through trans-splicing, and that this RNA fusion may be a prerequisite for cancer-causing chromosomal translocations (*23*). While this hypothesis had begun to implicate gene interactions in some mutations relevant to disease etiology, it had not involved genetic interactions in mutational mechanisms of evolutionary change. In contrast, the used-together-fused-together hypothesis offers a scientific perspective explaining why there are recurrent gene fusions both in evolution on the one hand (*24, 25*) and in genetic disease and cancer on the other (*26, 27*).

Here, we used bioinformatic techniques to test the used-together-fused-together (henceforth “used-fused”) hypothesis: whether genes that are used together are more likely to undergo a fusion mutation than random pairs of genes. While gene fusion in one species has been used to predict interactions in other species in bacteria and archaea (*18, 19*), here we examined relevant data from humans and other eukaryotes. More importantly, we attempted to distinguish the used-fused hypothesis from an explanation based on random mutation and natural selection alone: namely, that no mechanistic mutational reasons are involved, but rather genes that are used together, once they happen to undergo a random fusion mutation (specifically, a mutation in which mechanistic reasons relating to their being used together are not involved), produce a gene fusion that is then more likely to be favored by selection in comparison to the fusion of random gene pairs. Such differences in selective value could be due to bringing interacting gene products together more effectively or due to the more general possibility that genes already working together are relatively more predisposed to producing a beneficial fusion. Convergent evidence from the tests described below provides support for the used-fused hypothesis over the last and other hypotheses.

## Results

STRING is a large database providing information on interactions between proteins as well as protein fusions in numerous species (*28*). We used this database to identify pairs of separate human genes whose homologs are fused in other species (see Materials and Methods). We then compared the identified fusion-related gene pairs to a list of randomly generated gene pairs from the human genome (the control group; see Materials and Methods and brief explanation below) in order to test whether pair members of the fusion-related group tend to interact more with each other in humans than pair members of the control group. We used several metrics to investigate interactions between pair members: co-expression, co-localization of the pair members’ gene products in the cell, the tendency of the pair members to be found in the same topologically associating domain (TAD) in the genome and semantic similarity of their associated GO terms.

We generated the above-mentioned control group in two different ways, both controlling for distance between pair members as discussed shortly. First, we paired up randomly chosen protein-coding genes from the human genome (henceforth “genomic control”). Second, we draw at random gene pairs from the subset of STRING gene pairs not thought to have undergone fusion in other species (henceforth “STRING control”). For the purpose of testing the used-fused hypothesis, the last control is a conservative one, because gene pairs appear on STRING if they have already demonstrated one indicator of interaction or another (fusion being one such indicator). Since the distance between pair members is expected to be correlated with their co-functioning, in the process of generating the control groups we compiled lists of randomly chosen gene pairs in such manner that each distance between pair members observed in the fusion-related group was equally represented percentage-wise in the fusion-related and control groups (where distance is measured in terms of the number of coding genes separating between pair members), thus ensuring that all comparisons described below are controlled for the distances between pair members.

### Fusion-related pair members are more highly co-expressed than members of random pairs of the same intra-pair distance

Co-expression data for the human genome was downloaded from CoexpressDB (*29*) and consisted of four databases based on different co-expression platforms (Materials and Methods). For both the genomic and STRING controls, the co-expression of pair members was significantly higher in the fusion-related pairs than in the control pairs, for both the same-chromosome (p < 2.20E-16 in both the genomic and STRING control cases; one-sided Mann-Whitney (MW) test) and different-chromosome (p < 2.20E-16, genomic control and p < 3.00E-02, STRING control; one-sided MW test) groups (Table 1).

**Table 1a.** Co-expression comparisons between the fusion-related and control gene pairs, using the COXPRESSdb database Hsa-m2-v18-09

| Distance<sup>a</sup> | Genomic control |  |  | String control |  |  |
| --- | --- | --- | --- | --- | --- | --- |
|  | Group size<sup>b</sup> | p-value<sup>c</sup> | W<sup>c</sup> | Group size<sup>b</sup> | p-value<sup>c</sup> | W<sup>c</sup> |
| All pairs | 10438-103751 | <2.20E-16 | 3.74E+08 | 10438-102877 | <2.20E-16 | 4.67E+08 |
| Same chromosome | 2573-25101 | <2.20E-16 | 1.58E+07 | 2573-24227 | <2.20E-16 | 1.94E+07 |
| SC_0 | 745-6821 | <2.20E-16 | 1.98E+06 | 745-5963 | 9.33E-04 | 2.07E+06 |
| SC_1-100 | 1413-14130 | <2.20E-16 | 2.69E+06 | 1413-14130 | <2.20E-16 | 3.83E+06 |
| SC_100-500 | 269-2690 | 1.87E-10 | 2.78E+05 | 269-2690 | 4.98E-01 | 3.62E+05 |
| SC_500+ | 146-1460 | 2.62E-06 | 8.22E+04 | 146-1444 | 6.28E-01 | 1.07E+05 |
| Different chromosomes | 7865-78650 | <2.20E-16 | 2.25E+08 | 7865-78650 | <2.20E-16 | 2.89E+08 |

**Table 1b.** Co-expression comparisons between the fusion-related and control gene pairs, using the COXPRESSdb database Hsa-m-v18-10

| Distance<sup>a</sup> | Genomic control |  |  | String control |  |  |
| --- | --- | --- | --- | --- | --- | --- |
|  | Group size<sup>b</sup> | p-value<sup>c</sup> | W<sup>c</sup> | Group size<sup>b</sup> | p-value<sup>c</sup> | W<sup>c</sup> |
| All pairs | 9453-93533 | <2.20E-16 | 3.27E+08 | 9453-92866 | <2.20E-16 | 4.06E+08 |
| Same chromosome | 2279-21793 | <2.20E-16 | 1.34E+07 | 2279-21126 | <2.20E-16 | 1.66E+07 |
| SC_0 | 718-6183 | <2.20E-16 | 1.68E+06 | 718-5532 | 7.99E-05 | 1.81E+06 |
| SC_1-100 | 1182-11820 | <2.20E-16 | 2.51E+06 | 1182-11820 | <2.20E-16 | 3.51E+06 |
| SC_100-500 | 259-2590 | <2.20E-16 | 2.25E+05 | 259-2590 | 3.34E-02 | 3.12E+05 |
| SC_500+ | 120-1200 | 6.31E-05 | 5.67E+04 | 120-1184 | 6.60E-01 | 7.27E+04 |
| Different chromosomes | 7174-71740 | <2.20E-16 | 1.98E+08 | 7174-71740 | 2.53E-02 | 2.54E+08 |

**Table 1c.** Co-expression comparisons between the fusion-related and control gene pairs, using the COXPRESSdb database Hsa-r-v18-12

| Distance<sup>a</sup> | Genomic control |  |  | String control |  |  |
| --- | --- | --- | --- | --- | --- | --- |
|  | Group size<sup>b</sup> | p-value<sup>c</sup> | W<sup>c</sup> | Group size<sup>b</sup> | p-value<sup>c</sup> | W<sup>c</sup> |
| All pairs | 9336-92258 | <2.20E-16 | 2.81E+08 | 9336-91521 | <2.20E-16 | 3.65E+08 |
| Same chromosome | 2292-21818 | <2.20E-16 | 1.06E+07 | 2292-21081 | <2.20E-16 | 1.36E+07 |
| SC_0 | 687-5768 | <2.20E-16 | 1.51E+06 | 687-5048 | 2.90E-04 | 1.59E+06 |
| SC_1-100 | 1239-12390 | <2.20E-16 | 1.59E+06 | 1239-12390 | <2.20E-16 | 2.43E+06 |
| SC_100-500 | 241-2410 | <2.20E-16 | 1.92E+05 | 241-2410 | 2.63E-02 | 2.68E+05 |
| SC_500+ | 125-1250 | 5.35E-07 | 5.75E+04 | 125-1233 | 4.52E-01 | 7.66E+04 |
| Different chromosomes | 7044-70440 | <2.20E-16 | 1.70E+08 | 7044-70440 | <2.20E-16 | 2.31E+08 |

**Table 1d.**
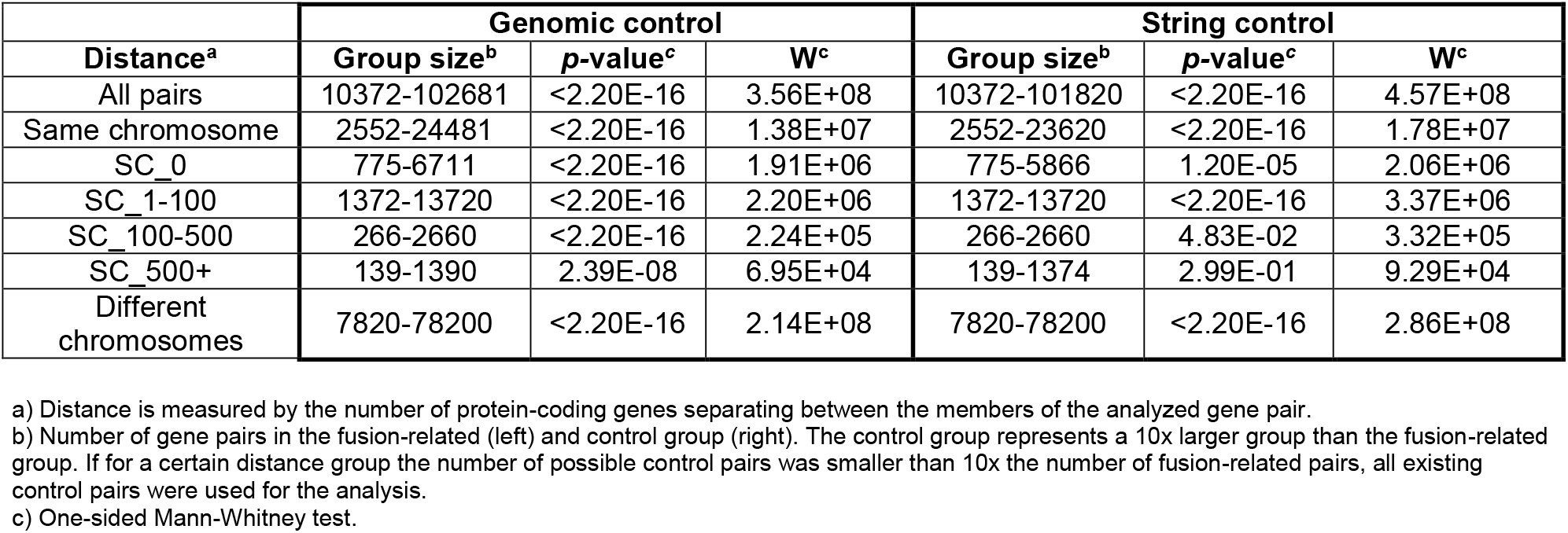
Co-expression comparisons between the fusion-related and control gene pairs using the COXPRESSdb database Hsa-u-v18-12

| Distance<sup>a</sup> | Genomic control |  |  | String control |  |  |
| --- | --- | --- | --- | --- | --- | --- |
|  | Group size<sup>b</sup> | p-value<sup>c</sup> | W<sup>c</sup> | Group size<sup>b</sup> | p-value<sup>c</sup> | W<sup>c</sup> |
| All pairs | 10372-102681 | <2.20E-16 | 3.56E+08 | 10372-101820 | <2.20E-16 | 4.57E+08 |
| Same chromosome | 2552-24481 | <2.20E-16 | 1.38E+07 | 2552-23620 | <2.20E-16 | 1.78E+07 |
| SC_0 | 775-6711 | <2.20E-16 | 1.91E+06 | 775-5866 | 1.20E-05 | 2.06E+06 |
| SC_1-100 | 1372-13720 | <2.20E-16 | 2.20E+06 | 1372-13720 | <2.20E-16 | 3.37E+06 |
| SC_100-500 | 266-2660 | <2.20E-16 | 2.24E+05 | 266-2660 | 4.83E-02 | 3.32E+05 |
| SC_500+ | 139-1390 | 2.39E-08 | 6.95E+04 | 139-1374 | 2.99E-01 | 9.29E+04 |
| Different chromosomes | 7820-78200 | <2.20E-16 | 2.14E+08 | 7820-78200 | <2.20E-16 | 2.86E+08 |

a) Distance is measured by the number of protein-coding genes separating between the members of the analyzed gene pair.
b) Number of gene pairs in the fusion-related (left) and control group (right). The control group represents a 10x larger group than the fusion-related group. If for a certain distance group the number of possible control pairs was smaller than 10x the number of fusion-related pairs, all existing control pairs were used for the analysis.
c) One-sided Mann-Whitney test.

We next divided the same-chromosome group into four separate sub-groups for a more detailed analysis: 1) SC_0: gene pairs whose pair members have no coding genes separating them; i.e., the pair members are neighbors; 2) SC_1-100: pair members are separated by 1 to 100 coding genes; 3) SC_100-500: pair members are separated by 100 to 500 coding genes; 4) SC_500+: pair members are separated by 500 or more coding genes.

Analysis revealed that the differences between fusion-related and control pairs were significant for the SC_0 (p < 2.20E-16 and p < 9.50E-04, genomic and STRING respectively; one-sided MW test) and SC_1-100 groups (p < 2.20E-16 in both the genomic and STRING control cases; one-sided MW test). For the SC_100-500 group, the results were significant for the genomic control case (p < 2.00E-10; one-sided MW test) in all co-expression databases, and for the STRING control in three of four databases (p < 4.90E-02; one-sided MW test). For the SC_500+ group, the results were significant for the genomic control only (p < 6.50E-05; one-sided MW test) and not for the STRING control.

To summarize, both for the same-chromosome and different-chromosome groups, we observe a general tendency whereby fusion-related pair members are more highly co-expressed than control pair members while controlling for distance between pair members. In the same-chromosome group, the significance of this comparison is larger for pairs whose members are closer to each other, although this effect may be due to the group sizes rather than the distance between pair members *per se*, as we obtain strongly significant results also for different-chromosome pair members.

### Co-localization is higher for fusion-related pair members than for members of random pairs of the same intra-pair distance

Next, we compared the extent to which the protein products of pair members localize to the same cellular compartment between the STRING fusion-related gene pairs and the controls using a database of protein sub-cellular localizations downloaded from the CoexpressDB website (*29, 30*). For both the same-chromosome and different-chromosome groups, the co-localization of pair members’ gene products was significantly higher in the fusion-related than in the control pairs (p < 2.20E-16 in both the genomic and STRING control cases; one-sided Fisher exact test) (Table 2).

**Table 2.** Co-localization comparisons between the fusion-related and control gene pairs, using the Psort database from COXPRESSdb portal

| Distance<sup>a</sup> | Genomic control |  |  |  | String control |  |  |  |
| --- | --- | --- | --- | --- | --- | --- | --- | --- |
|  | Group size<sup>b</sup> | Number of positives<sup>c</sup> | p-value<sup>d</sup> | Odds ratio<sup>d</sup> | Group size<sup>b</sup> | Number of positives<sup>c</sup> | p-value<sup>d</sup> | Odds ratio<sup>d</sup> |
| All pairs | 10747-106502 | 7344-50966 | <2.20E-16 | 2.352 | 10747-105819 | 7344-62898 | <2.20E-16 | 1.473 |
| Same chromosome | 2644-25472 | 1979-12660 | <2.20E-16 | 3.012 | 2644-24789 | 1979-14316 | <2.20E-16 | 2.177 |
| SC_0 | 760-6632 | 436-3585 | 4.46E-02 | 1.144 | 760-5966 | 436-3430 | 5.42E-01 | 0.995 |
| SC_1-100 | 1457-14570 | 1246-7043 | <2.20E-16 | 6.310 | 1457-14570 | 1246-8397 | <2.20E-16 | 4.341 |
| SC_100-500 | 272-2720 | 172-1309 | 1.23E-06 | 1.854 | 272-2720 | 172-1575 | 5.03E-02 | 1.250 |
| SC_500+ | 155-1550 | 125-723 | <2.20E-16 | 4.762 | 155-1533 | 125-914 | 7.28E-08 | 2.820 |
| Different chromosomes | 8103-81030 | 5365-38306 | <2.20E-16 | 2.185 | 8103-81030 | 5365-48582 | <2.20E-16 | 1.309 |

- a) Distance is measured by the number of protein-coding genes separating between the members of the analyzed gene pair. b) Number of gene pairs in the fusion-related (left) and control group (right). The control group represents a 10x larger group than the fusion-related group. If for a certain distance group the number of possible control pairs was smaller than 10x the number of fusion-related pairs, all existing control pairs were used for the analysis. c) Number of co-localized pair members from the fusion-related group (left) and the control group (right). d) One-sided Fisher exact test.

Further analysis of the same-chromosome group revealed significant differences between fusion-related gene pairs and controls in the SC_1-100 (p < 2.20E-16 in both the genomic and STRING control cases; one-sided Fisher exact test) and SC_500+ (p < 2.20E-16 and p < 7.30E-08 in the genomic and STRING controls, resp.; one-sided Fisher exact test) groups. For neighboring pair members (SC_0) and the SC_100-500 group, significant results were obtained only in the genomic control case (p = 4.46E-02 and p = 1.23E-06, respectively; one-sided Fisher exact test). Additionally, for the SC_100-500 group, the STRING control results were marginally significant (p = 5.03E-02; one-sided Fisher exact test)

To summarize, co-localization of pair members’ gene products is in general significantly higher for fusion-related than control pairs, both for the same-chromosome and different-chromosome groups.

### Fusion-related pair members are more often found in the same TAD than members of random pairs of the same intra-pair distance

A topologically associating domain (TAD) is a region in the genome whose DNA sequences interact physically preferentially with each other and are found in close proximity to each other in 3D due to the DNA’s 3D structure. We used a database of TAD coordinates within the human genome downloaded from the 3D Genome Browser website (*31*), to investigate whether fusion-related pair members tend to be found in the same TAD more frequently than the members of random pairs. Importantly, note that, as in previous comparisons, the 2D distance between pair members was controlled for by matching the distances between the fusion-related and control groups. We analyzed only same-chromosome pair members since the database does not provide information on interactions across different chromosomes. Examining all of the same-chromosome gene pairs together, we found that fusion-related members are more often found in the same TAD than control pair members (p < 2.20E-16 in both the genomic and STRING control cases; one-sided MW test) (Table 3). Further analysis of the same-chromosome gene pairs revealed that these differences are mainly driven by the SC_1-100 group (p < 2.20E-16 in both the genomic and STRING control cases; one-sided MW test). Smaller yet still significant differences exist in same-TAD presence between neighboring pair members (SC_0) and control pairs (p = 2.04E-02 and p = 2.96E-03, genomic and STRING controls, resp.; one-sided MW test). Although small, this effect is impressive given that neighboring genes are likely to be present in the same TAD due to their proximity to each other alone, thus reducing the potential for finding a difference in same-TAD-presence between the groups to begin with.

**Table 3.** Same-TAD presence comparisons between the fusion-related and control gene pairs

| **Distance<sup>a</sup>** | **Genomic control** |  |  | **String control** |  |  |
| --- | --- | --- | --- | --- | --- | --- |
|  | **Group size<sup>b</sup>** | ***p*-value<sup>c</sup>** | **W<sup>c</sup>** | **Group size<sup>b</sup>** | ***p*-value<sup>c</sup>** | **W<sup>c</sup>** |
| All pairs | 2747-27087 | <2.20E-16 | 4.64E+07 | 2747-25686 | <2.20E-16 | 4.40E+07 |
| Same chromosome | 2747-27087 | <2.20E-16 | 4.64E+07 | 2747-25686 | <2.20E-16 | 4.40E+07 |
| SC_0 | 806-7677 | 2.04E-02 | 3.23E+06 | 806-6292 | 2.96E-03 | 2.69E+06 |
| SC_1-100 | 1494-14940 | <2.20E-16 | 1.64E+07 | 1494-14940 | <2.20E-16 | 1.58E+07 |
| SC_100-500 | 281-2810 | 2.10E-01 | 3.97E+05 | 281-2810 | 2.08E-01 | 3.97E+05 |
| SC_500+ | 166-1660 | 1.00E+00 | 1.38E+05 | 166-1644 | 1.00E+00 | 1.36E+05 |

a) Distance is measured by the number of protein-coding genes separating between the members of the analyzed gene pair.
b) Number of gene pairs in the fusion-related (left) and control group (right). The control group represents a 10x larger group than the fusion-related group. If for a certain distance group the number of possible control pairs was smaller than 10x the number of fusion-related pairs, all existing control pairs were used for the analysis.
c) One-sided Mann-Whitney test.

For pairs whose members are farther from each other, namely for the SC_100-500 and SC_500+ groups, no significant differences were found between the fusion-related and control pairs (genomic and STRING). Likely in these groups the pair members are too far apart to be present in the same TAD in most if not all cases.

### GO terms are more similar between fusion-related pair members than between members of random pairs of the same intra-pair distance

Finally, we compared the semantic similarity of the GO terms associated with pair members in the fusion-related and control groups using GOGO (*32*). We performed this analysis separately for each of the three main GO categories: Biological Process (BP), Molecular Function (MF) and Cellular Component (CC). Compared to the control groups, we found a significantly higher semantic similarity of GO terms when examining the entire group of same-chromosome pair members (p < 2.20E-16, genomic and STRING controls resp.; one-sided MW test), as well as the group of different-chromosome pair members (p < 2.20E-16 and p < 3.40E-06, for genomic and STRING controls; one-sided MW test) for all three main GO categories: BP, MF and CC (Table 4).

**Table 4a.** GO term BP (biological process) semantic similarity comparisons between the fusion-related and control gene pairs

| Distance<sup>a</sup> | Genomic control |  |  | String control |  |  |
| --- | --- | --- | --- | --- | --- | --- |
|  | Group size<sup>b</sup> | p-value<sup>c</sup> | W<sup>c</sup> | Group size<sup>b</sup> | p-value<sup>c</sup> | W<sup>c</sup> |
| All pairs | 10389-102983 | <2.20E-16 | 7.48E+08 | 10389-102329 | <2.20E-16 | 5.83E+08 |
| Same chromosome | 2459-23683 | <2.20E-16 | 4.23E+07 | 2459-23029 | <2.20E-16 | 3.64E+07 |
| SC_0 | 662-5713 | 4.92E-01 | 1.89E+06 | 662-5076 | 1.00E+00 | 1.54E+06 |
| SC_1-100 | 1379-13790 | <2.20E-16 | 1.61E+07 | 1379-13790 | <2.20E-16 | 1.45E+07 |
| SC_100-500 | 261-2610 | <2.20E-16 | 4.82E+05 | 261-2610 | 1.07E-01 | 3.56E+05 |
| SC_500+ | 157-1570 | <2.20E-16 | 1.72E+05 | 157-1553 | 2.22E-02 | 1.34E+05 |
| Different chromosomes | 7930-79300 | <2.20E-16 | 4.34E+08 | 7930-79300 | 3.38E-06 | 3.24E+08 |

**Table 4b.** GO term MF (molecular function) semantic similarity comparisons between the fusion-related and control gene pairs

| Distance<sup>a</sup> | Genomic control |  |  | String control |  |  |
| --- | --- | --- | --- | --- | --- | --- |
|  | Group size<sup>b</sup> | p-value<sup>c</sup> | W<sup>c</sup> | Group size<sup>b</sup> | p-value<sup>c</sup> | W<sup>c</sup> |
| All pairs | 10491-103850 | <2.20E-16 | 7.07E+08 | 10491-103167 | <2.20E-16 | 6.01E+08 |
| Same chromosome | 2473-23670 | <2.20E-16 | 4.10E+07 | 2473-22987 | <2.20E-16 | 3.71E+07 |
| SC_0 | 675-5690 | 6.55E-01 | 1.90E+06 | 675-5025 | 9.97E-01 | 1.58E+06 |
| SC_1-100 | 1379-13790 | <2.20E-16 | 1.55E+07 | 1379-13790 | <2.20E-16 | 1.48E+07 |
| SC_100-500 | 261-2610 | 4.66E-09 | 4.14E+05 | 261-2610 | 4.71E-01 | 3.42E+05 |
| SC_500+ | 158-1580 | <2.20E-16 | 1.77E+05 | 158-1562 | 3.53E-06 | 1.50E+05 |
| Different chromosomes | 8018-80180 | <2.20E-16 | 4.06E+08 | 8018-80180 | 9.08E-11 | 3.35E+08 |

**Table 4c.** GO term CC (cellular component) semantic similarity comparisons between the fusion-related and control gene pairs

| Distance<sup>a</sup> | Genomic control |  |  | String control |  |  |
| --- | --- | --- | --- | --- | --- | --- |
|  | Group size<sup>b</sup> | p-value<sup>c</sup> | W<sup>c</sup> | Group size<sup>b</sup> | p-value<sup>c</sup> | W<sup>c</sup> |
| All pairs | 10662-105805 | <2.20E-16 | 8.09E+08 | 10662-104955 | <2.20E-16 | 6.65E+08 |
| Same chromosome | 2574-24925 | <2.20E-16 | 4.63E+07 | 2574-24075 | <2.20E-16 | 4.05E+07 |
| SC_0 | 723-6415 | 6.56E-04 | 2.49E+06 | 723-5582 | 7.32E-01 | 1.99E+06 |
| SC_1-100 | 1426-14260 | <2.20E-16 | 1.68E+07 | 1426-14260 | <2.20E-16 | 1.56E+07 |
| SC_100-500 | 265-2650 | <2.20E-16 | 4.71E+05 | 265-2650 | 1.14E-01 | 3.67E+05 |
| SC_500+ | 160-1600 | <2.20E-16 | 1.83E+05 | 160-1583 | 2.95E-05 | 1.51E+05 |
| Different chromosomes | 8088-80880 | <2.20E-16 | 4.68E+08 | 8088-80880 | <2.20E-16 | 3.75E+08 |

a) Distance is measured by the number of protein-coding genes separating between the members of the analyzed gene pair.
b) Number of gene pairs in the fusion-related (left) and control group (right). The control group represents a 10x larger group than the fusion-related group. If for a certain distance group the number of possible control pairs was smaller than 10x the number of fusion-related pairs, all existing control pairs were used for the analysis.
c) One-sided Mann-Whitney test.

Further analysis of the same-chromosome group revealed results similar to those of the co-localization analysis. Significant differences between the fusion-related and control pairs were found for all three main GO categories in the SC_1-100 (p < 2.20E-16, genomic and STRING controls resp.; one-sided MW test) and SC_500+ (p < 2.20E-16 and p < 2.30E-02, genomic and STRING controls resp.; one-sided MW test) groups. For the SC_0 group, significant differences were found only in the cellular component GO category in the genomic control case (p = 6.56E-04, ; one-sided MW test) but not in the STRING control case (p = 7.32E-01, ; one-sided MW test). For the SC_100-500 group, significant results were obtained only in the genomic control case, in all three GO categories: BP, MF and CC (p < 4.70E-09; one-sided MW test).

To summarize, the semantic similarity between the GO terms associated with pair members is generally significantly higher in the fusion-related than control pairs, both for same- chromosome and different-chromosome pair members. Further analysis reveals that for pair members on the same chromosome, these differences tend to be found in more distant pair members, but not in neighbors, possibly because neighboring genes are more likely in general to share the same GO category.

### Fusion-related gene pairs are more highly represented than random gene pairs in the list of human cancer gene fusions

As will be discussed further, and while recognizing the possibility that the events considered may include not only fusion but also fission (see Discussion), the results so far are consistent with the possibility that genes that interact more with each other are more likely to become fused in the course of evolution. However, as will be discussed further, these results leave open the possibility that interacting genes, once fused by a random mutation, generate a fusion that is more likely to be favored by selection in comparison to the fusion of a random pair of genes— to distinguish from the possibility that interacting genes are more likely to undergo a fusion mutation, as suggested by the TRIMCyp starting example and the used-fused hypothesis. To distinguish between these possibilities, we compared the STRING fusion-related gene pairs to the control pairs in terms of their presence in a large database of gene pairs that undergo fusion in human cancers. It is unlikely that selection, following random mutation, would favor the fusion of the same pairs of genes in both human cancerous cells and non-human organisms, given the vastly different selection pressures involved in these two cases. In contrast, if it is genes that interact that can undergo a fusion mutation, then a far smaller set of gene pairs can potentially fuse either in evolution or in cancer, enabling an overlap between evolutionary and cancer fusions despite the vastly different selection pressures involved. Therefore, a statistically significant overlap between the evolutionary-fusion–related gene pairs of STRING and gene pairs that are fused in cancer would favor a mutational-over a strictly selection-based hypothesis.

To test for a significant overlap between the evolutionary and cancer fusions we used cancer fusion data downloaded from the Fusion-GDB and COSMIC portals (*33–35*). Results show that cancer databases are enriched in same-chromosome evolutionary-fusion–related gene pairs compared to control pairs, both in the genomic (p = 5.07E-11; one-sided Fisher exact test) and STRING (p = 2.47E-04; one-sided Fisher exact test) control cases (Table 5). Further analysis of the same-chromosome group revealed that this significance is driven by neighboring (SC_0) pair members (p = 2.83E-12 and p = 9.80E-05, genomic and STRING controls, resp.; one-sided Fisher exact test). For pair members more distant from each other (SC_1-100, SC_100-500 and SC_500+), the overlap with the cancer-fusion list was not significantly larger in the evolutionary-fusion–related pairs than in the control pairs. Finally, for the different-chromosome group, the overlap was significantly larger in the evolutionary-fusion–related than in the control pairs in the genomic control case (p = 5.40E-03, one-sided Fisher exact test) but not in the STRING control case.

**Table 5.** Comparisons between the fusion-related and control gene pairs in terms of their presence on cancer fusion database

| Distance<sup>a</sup> | Genomic control |  |  |  | String control |  |  |  |
| --- | --- | --- | --- | --- | --- | --- | --- | --- |
|  | Group size<sup>b</sup> | Number of positives<sup>c</sup> | p-value<sup>d</sup> | Odds ratio<sup>d</sup> | Group size<sup>b</sup> | Number of positives<sup>c</sup> | p-value<sup>d</sup> | Odds ratio<sup>d</sup> |
| All pairs | 11293-112930 | 168-919 | 1.02E-11 | 1.841 | 11293-111386 | 168-1174 | 3.24E-05 | 1.418 |
| Same chromosome | 2762-27620 | 162-906 | 5.07E-11 | 1.837 | 2762-26076 | 162-1134 | 2.47E-04 | 1.370 |
| SC_0 | 817-8170 | 126-630 | 2.83E-12 | 2.182 | 817-6640 | 126-718 | 9.80E-05 | 1.504 |
| SC_1-100 | 1498-14980 | 35-271 | 9.29E-02 | 1.298 | 1498-14980 | 35-413 | 8.51E-01 | 0.844 |
| SC_100-500 | 280-2800 | 0-5 | 1.00E+00 | 0.000 | 280-2800 | 0-3 | 1.00E+00 | 0.000 |
| SC_500+ | 167-1670 | 1-0 | 9.09E-02 | Inf | 167-1656 | 1-0 | 9.16E-02 | Inf |
| Different chromosomes | 8531-85310 | 6-13 | 5.40E-03 | 4.618 | 8531-85310 | 6-40 | 2.37E-01 | 1.500 |

a) Distance is measured by the number of protein-coding genes separating between the members of the analyzed gene pair.
b) Number of gene pairs in the fusion-related (left) and control group (right). The control group represents a 10x larger group than the fusion-related group. If for a certain distance group the number of possible control pairs was smaller than 10x the number of fusion-related pairs, all existing control pairs were used for the analysis.
c) Number of pairs from the fusion-related (left) and control (right) groups that have been observed to fuse in human cancers.
d) One-sided Fisher exact test.

In the same chromosome group, the number of gene pairs overlapping with the cancer fusion database decreased with increasing distance between pair members, again likely due to decreasing group sizes, the largest percent of overlapping pairs being in the SC_0 group (15.42%), followed by the SC_1-100 group (2.34%). The SC_100-500 group showed no overlap and the SC_500+ group had only one overlapping pair. In the different-chromosome group, six overlapping pairs were found (0.07%)

## Discussion

We first found that many of the pairs of genes that are fused in other organisms but separate in humans are neighboring genes in humans. Under the assumption that these cases represent fusions primarily rather than fissions (*36*) (see more below), this raises the possibility that most fusions are generated by random transcriptional read-through or random deletion mutations that connect neighboring or proximal genes in a single random mutational event, and thus the results may be explained as follows: It is known from past literature that nearby genes are more likely to be working together than genes remote from each other (*37–40*). If they are so due to reasons unrelated to fusion, then the results may be correlational in a manner understandable from random mutation, since only nearby genes can be fused by random read-through or deletion mutations (*H*_1_).

However, our findings show that this explanation is not sufficient on its own. First, genes fused in other organisms are more likely to be working together in humans even when controlling for distance between pair members. Thus, even among pairs of neighboring genes of the same distance between the pair members, those pairs that work together more are more likely to become fused. Various measures of “working together” provide cross validation for this finding, including co-expression, co-localization, same-TAD presence and semantic similarity of GO terms. However, *H*_1_ does not predict a difference in fusion propensity between neighbors that work together less or more.

Second, the random read-through- or deletion-mutation hypothesis does not directly account for the fact that the used-fused effect exists also in pair members that are distant from each other, whether in the same or in different chromosomes (Tables 1, 2 and 4). While one could hypothetically argue that all fused genes were neighbors at the moment of fusion in the species where they were fused, this is a restrictive assumption. Indeed, the TRIMCyp case mentioned in the introduction is just one example of fusion of non-neighbors that is difficult to explain by random mutation due to its recurrence. Furthermore, the overlap between the evolutionary and cancer fusions includes several gene pairs whose members are distant from each other in humans yet become fused from a distance in cancer. Third, the finding that genes in the same TAD are more likely to undergo a fusion mutation compared to genes from different TADs while controlling for distance between pair members cannot be well explained by read-through or deletion mutations.

An alternative hypothesis based on random mutation with which to account for the results is that by random mutation, genes become fused, and of these random gene fusions, the ones whose components previously interacted are more likely to be favored by selection because their history of having worked together makes them predisposed to perform better once fused relative to pairs that have not already worked together (*H*_2_). This hypothesis admits any type of random mutation, including but not limited to read-through and deletion mutations connecting nearby genes, and thus in principle could account for observed fusions of pair members that work together both that are nearby and that are remote from each other, as well as for the increased fusion tendency of neighboring genes that interact more tightly.

However, this hypothesis (*H*_2_) does not account for the cancer-overlap result: the result that genes that were fused in other organisms in the course of evolution are more likely than random pairs to be fused in human cancers. Notice that selection in other organisms (e.g., in primates) favors mutations that increase survival and reproduction at the organismal level (e.g., improving tree-climbing abilities or digestion of relevant food sources or innumerable other qualities), whereas “selection” among human cancer cells favors mutations that increase the ability of the cell to proliferate as a cancerous cell and thus have an increased chance of being observed in cancer samples, which is not expected to match organismal survival and reproduction systematically and often comes at the expense of the latter. This contrast between the fundamental nature of the selection pressures involved leaves unexplained why there would be a statistically significant overlap between the list of evolutionary fusions in other organisms and the list of human cancer fusions under the purely selection-based explanation *H*_2_.

In contrast, all of the findings are consistent with the used-together-fused-together hypothesis—the hypothesis that genes that are used together are more likely to become fused for mechanistic mutational reasons inherent to their interaction (*13,14*). First, since neighboring genes are much more likely to be working together than non-neighboring genes (*37–40*), this hypothesis is immediately consistent with the fact that many fusions observed are between neighbors. Second, unlike *H*_1_, it immediately accounts for the findings that, among same-distance neighboring genes, those that interact more closely with each other are more likely to be fused; that genes in the same TAD are more likely to be fused than random same-distance pairs; and that distant genes that work together are more likely than random same-distance pairs to fuse. Third, unlike *H*_2_, the effect whereby genes that work together are more likely to undergo a fusion mutation could account for the propensity of the same gene pairs to undergo fusion in both evolution and cancer without hindrance. The resulting fusion mutations could then undergo different selection pressures in each case, leaving a small but statistically significant overlap between evolutionary and cancer fusions, as observed. In other words, mutation is a primary factor limiting the set of gene pairs with fusion potential, explaining the cancer-evolution overlap.

The cancer-evolution overlap also addresses the possibility that some or all of the cases observed represent fission rather than fusion events and therefore do not support the used-fused hypothesis. We know that in the case of cancer, these are indeed fusions, not fissions. Therefore, it would be unusual if the gene pairs that repeat in cancer and evolution are fusion events in cancer but fission events in evolution. That would seem to suggest that genes were once fused, then separated in humans and some other organisms, and are now fusing together again in human cancers. To avoid the used-fused hypothesis here one would have to argue that these genes were separated by random mutation and natural selection in humans and some other organisms, and are now fused back by random mutation and “selection” for survival and proliferation in human cancer tissues. It is more parsimonious to suggest that these pairs were fused both in evolution and in cancer, in both cases due to the same mutational mechanisms.

Besides accounting for the evolution-cancer fusions overlap, the used-fused hypothesis also offers a more parsimonious explanation than *H*_2_ for the other results above-mentioned. To avoid the used-fused hypothesis while using the arsenal of alternatives considered here, one must invoke *H*_2_ to explain the finding that genes that are co-expressed more are more likely to become fused also when they are distant from each other (assuming indeed that not only neighboring genes become fused, consistent with the TRIMCyp study case, Tables 1–4 and the cancer-overlap cases). However, it would be highly counterintuitive to use that same hypothesis, *H*_2_, to account for the fact that many fused genes are neighbors: that would require ignoring the obvious potential of neighbors to be fused more often than non-neighbors for mutational reasons (even if such mutational reasons are limited to random read-through or deletion mutations). However, adding *H*_1_ to account for the fact that many fusion-related genes are neighbors means using two different hypotheses to explain even only this limited part of the results.

Again unless making the problematic assumptions that all fused genes were neighbors prior to fusing, or that all of the findings pertain to cases of fission, adding the TADs result would only make avoiding the used-fused hypothesis more complicated: We will now have to add another hypothesis, *H′*_1_, that genes in the same TAD are more likely than others to be working together, and, due to their proximity in 3D but unrelated to the fact that they work together, are also more likely to undergo a fusion mutation. However, in comparison to *H*_1_, it is harder to make a clear separation here between the points *a*) that genes that work together are more likely than otherwise to be in the same TAD, and *b*) that genes that are in the same TAD are more likely to undergo a fusion mutation, as to then argue that those are separate, unrelated points that coincidentally overlap for involving TADs. Such a separation is counterintuitive because the same mechanisms due to which being in the same TAD facilitates genetic interaction are also expected to facilitate fusion mutations, as argued by the used-fused hypothesis: genes that work together are more likely to be expressed and thus have their chromatin open at the same time and place in the nucleus, allowing for various downstream mechanisms, whether retroposition, trans-splicing, recombination or more to increase the chance of fusion (*13,14*). Thus, to explain the co-expression, co-localization, same-TAD presence and GO terms results without resorting to mutational mechanisms, not only one needs to invoke three different hypotheses, *H*_1_, *H′*_1_ and *H*_2_, when the mutational one (the used-fused hypothesis) agrees with all of the findings in one, but in addition one needs to make such a counterintuitive separation between points *a* and *b* above to distinguish *H′*_1_ from the used-fused hypothesis. It is in addition to this added parsimony of the used-fused hypothesis that the latter explains also the cancer-overlap results, which the other hypotheses do not.

Paralleling points *a* and *b* of *H′*_1_ above, *H*_1_ argues that *a*) neighbors are more likely than non-neighbors to be working together; and *b*) a read-through mutation can easily occur by chance; and that it is a coincidence that both points pertain to neighboring genes. However, the facts that the used-fused hypothesis is superior to *H′*_1_ in explaining the TADs result and that *H′*_1_ is similar to *H*_1_ enables viewing read-through and deletion mutations connecting nearby genes from another angle. While these mutations may appear easy to obtain at random, they actually involve a minimally mechanistic consideration: genome architecture endows these mutations with their potential effect (they also require a successful alternative splicing to follow, which has been taken for granted so far). Because the invocation of a mechanism here is minimal, it could be seen as fitting with the random mutation view in the absence of other data. However, the presence of the other findings obtained here raises the possibility that the used-fused framework explains gene fusions better; that additional mutational mechanisms besides read-through and deletion mutations may be involved in the fusion of neighbors; and that the read-through case is just an extreme example of this framework (extreme by involving a minimal mutational consideration).

In fact, given parsimony considerations, an extension of the used-fused hypothesis now suggests itself, namely that genes that are used together are incomparably more likely than others not only to be fused together by a fusion mutation, but also, when initially distant, to be moved by a translocation mutation to the same neighborhood. In fact, the same sorts of mechanisms proposed for the fusion case could be proposed for the case of moving to the same neighborhood. One may even expect that genes that interact from afar usually first move to the same neighborhood and only later in evolution become fused, where much evolutionary time may elapse between the steps of interacting from afar, translocating to the same neighborhood and fusing (*14, 25*). This extension of the used-fused hypothesis may help to explain why neighboring genes, or genes in the same TAD, are more likely than other genes to be working together in the first place. Absent such a mutational explanation, one has to either accept these facts as “just so” or invoke random mutation and natural selection (rm/ns) based arguments, such as that selection will favor the moving to the same neighborhood of genes that work together over those that do not because this will save energy or time or avoid errors in the process of their expression, and that selection will favor the fusion of such genes because it will save the time of bringing the products together—reasons that are questionable based on the minute economic considerations they invoke (*22*). In contrast, both phenomena of fusion and neighborhoods of collaborating genes can be accounted for by the extended used-fused hypothesis under one explanation and without resorting to such considerations.

In summary, the fact that genes that work together in one species, both neighboring and distant genes, are more likely than random pairs to be found fused in other species; the fact that neighbors are more commonly found fused in other species than remote genes; the fact that neighbors that interact more tightly are more likely to be found fused than neighbors that interact less tightly; the fact that genes in the same TAD are more likely to be found fused than genes of the same distance in different TADs; and the fact that the list of gene fusions in human cancers overlaps in a statistically significant manner with the list of evolutionary gene fusions in other species all lend themselves to the hypothesis that genes used together are fused together more than others for mutational mechanistic reasons.

## Implications

Thus far, investigators have had to rely on random mutation, natural selection, and random genetic drift to explain genome-organization evolution. However, that approach seemed to invoke either pure chance (e.g., ref. 41) or the reliance on minute economic considerations in order to explain the empirical patterns.

Here we found evidence for the used-fused hypothesis: the hypothesis that genes that are used together repeatedly and persistently in the course of evolution are more likely than others to undergo a fusion mutation. This hypothesis offers to account for the recurrence of gene fusions in both evolution on the one hand (*24, 25*) and in genetic disease and cancer on the other hand (*26, 27*) under one umbrella. We further hypothesized that mutational mechanisms are relevant not only to fusion but also to translocation of genes that interact to the same neighborhood, which would extend this umbrella to explain why neighboring genes and genes in the same TAD are more likely to interact than random pairs of genes (*37–40*). Thus, the extended used–fused hypothesis offers a way to avoid the paradox of minute economic considerations and exemplifies the possibility that mutational mechanisms are an important contributor to the evolution of genome organization.

This leads to an important point. Consistent with Gilbert’s hypothesis of genes in pieces (*42*), the intron-exon structure of eukaryotic genes allows the breakpoints of random rearrangement mutations to fall outside of exons, thus facilitating the evolutionary shuffling of whole exons without disrupting them and allowing for the generation of new combinations of exons (*42*). However, our results suggest that exon shuffling is not only the result of random mutation, and that the intron-exon structure may facilitate exon shuffling in a different way: they suggest that exons first interact from afar, and their interaction leads them mechanistically to be translocated to interact in cis and to become fused via mutational mechanisms (*14*). The contrast between exon shuffling by random mutations and the used-fused–driven shuffling is particularly clear in cases where the same exons are trans-spliced in one species or population and cis-spliced in another, as is the case of the exons of the eri-6 and eri-7 genes in *C. elegans* strain N2 and the corresponding exons of the fused homologs in *C. briggsae* (*43*), or in cases where some functions, such as the production of fatty acids from acetyl-CoA, are achieved by multiple single-module proteins in one taxon but by a single multi-module protein in another (*44*). Such evidence is not engaged by hypotheses of exon shuffling based on random mutation, but is consistent with the idea that mutational mechanisms, involving also the splicing machinery, play a role in exon shuffling (*14, 25*).

In addition to an effect on genome organization, another important consequence of the used-fused hypothesis is that mutational mechanisms could contribute to the observed fitness distribution of mutations. According to the general notion of random mutation—i.e., that mutation occurs as an accident to the genome due to physico-chemical reasons unrelated to the biology of the organism—it was proposed that detrimental mutations should be more common than beneficial ones (*45*). Only later was it discovered that the vast majority of substitutions appear neutral or nearly so (*46, 47*). But why should the vast majority of mutations be neutral? Some possibilities are synonymous mutations (*47*), that the majority of the genome is non-functional and thus the majority of mutations are of no effect (*48*), and that the majority of the genome consists of regulatory as opposed to coding regions and that mutations in the former may often have little effect (*49, 50*). However, not mutually exclusive with these possibilities, our results suggest that mutational mechanisms may also affect the fitness distribution of mutations: under the used-fused hypothesis, gene-fusion mutations are less accidental and therefore may be expected to be less disruptive to fitness as compared to random mutations. This raises the possibility that the fitness distribution of fusion mutations may have leaned more to the detrimental side than it does in reality if fusion mutations were purely random. Future studies may reveal further connections between mutational mechanisms and the empirical fitness distribution of mutations, not only for fusion mutations but for other mutations as well (*51*).

Another consequence of the used-fused hypothesis pertains to parallelism. It has been suggested that more closely related species are more likely to exhibit parallelism (e.g., refs. 52, 53) because they experience more similar selection pressures and have more similar genetic and developmental backgrounds based on which random mutations have their phenotypic effects (*54*). However, if mutational mechanisms and hence the state of the genome affect the probabilities of specific mutations, then mutational mechanisms potentially constitute an additional reason for this effect. If genes that interact tightly are more likely than others to undergo a fusion mutation, then multiple species that share this interaction may undergo the same or similar fusion/s independently, thus greatly increasing the probability of fusion parallelism in evolution in general and in adaptive evolution specifically. As shown by our starting example, the TRIMCyp fusion happened at least twice independently in two different simian lineages, and evidence exists showing that it provided added protection against certain lentiviruses (*4, 5*). It is hard to explain this recurrence by rm/ns alone. This conclusion is consistent with other recent empirical findings connecting mutational phenomena to parallelism in evolution in general and adaptive evolution specifically (*55–58*), such as the finding that the high rates of deletion of a specific enhancer are responsible for the parallel and likely adaptive loss of the pelvic hindfin in freshwater sticklebacks (*55*); and that the human hemoglobin S mutation, which protects against malaria in heterozygotes and causes sickle-cell anemia in homozygotes, originates significantly more rapidly than expected by chance for this mutation type, especially in Africans (*51*), strengthening the possibility that this mutation has after all arisen multiple independent times (*59*).

That mutational phenomena may enable extensive parallelism is relevant to the interpretation of phylogenetic evidence. If mutational mechanisms underlie gene fusion, then even when the majority of species in a monophyletic clade share a certain gene fusion, it could be that the common ancestor shared the tendency to generate the fusion, and that the fusion arose later independently multiple times. This new interpretation resolves a contradiction in previous data, where authors examining relatively more distant species concluded that fusions are more common than fissions (*36, 60*), and authors examining relatively more related species concluded the opposite (*61, 62*). If related species share a tendency to fuse the same pairs of genes, yet this possibility is ignored, then such contradictory results would indeed emerge. If, instead, this possibility is taken into account, then the data raises the possibility that fusions are always more common than fissions but often occur in parallel in related species.

Finally, consider the phenomenon of gene duplication via mutational mechanisms such as non-allelic homologous recombination, non-homologous end-joining, retroposition and others (*63–69*). It is hard to argue that it is because these mechanisms allowed for gene duplication that they evolved under rm/ns—such a benefit is a long-term one, whereas rm/ns is generally based on individual-level, immediate benefits (*70,71*); yet it is hardly possible to imagine evolution as we know it without the existence of these mechanisms, which are in fact of fundamental importance to evolution (*72, 73*). Likewise in the long-term, it is of interest to note that the chunking of pieces of information that are repeatedly used together into a single unit is a powerful principle across different processes of information acquisition (*74–77*) and that evolution has been thought of as such a process. As is the case with gene duplication, noting this potential benefit of fusing genes that work together is not to say that the used-fused effect itself evolved by rm/ns based on this benefit, but rather to recognize that it is an interesting and potentially important property of the genetic system as a whole, whose own origin requires further thinking.

## Acknowledgements

We thank Shahar Barbash for assistance with initial analyses, Eugene Koonin for suggestions and Kim Weaver for extensive help throughout the project.

## Funding

This publication was made possible through the support of a grant from the John Templeton Foundation. The opinions expressed in this publication are those of the authors and do not necessarily reflect the views of the John Templeton Foundation.

## Materials and Methods

### Identification of fusion-related gene pairs

We extracted from the STRING database (*28*) all pairs of interacting human proteins that have a non-zero fusion score, which indicates evidence that homologs of these proteins are fused in another or other species (henceforth “fusion-related”). While STRING provides a score indicating the level of confidence in the evidence of a particular type of interaction, in this case of a fusion we did not limit the analysis to only a subset of scores, in order to avoid missing true pairs of fusion candidates. Next, each pair of identified fusion-related proteins was mapped back to the genes that express them to create a list of fusion-related gene pairs. Since multiple protein products can be produced by the same gene(s), we scanned the resulting list for redundancy and removed repeating gene pairs, to ensure that each gene pair was represented only once in the list.

### Analysis of genomic distances between fusion-related gene pair members

The distance between genes might affect their tendency to interact. In order to control for this potential effect, we grouped the analyzed gene pairs by the distance between pair members. The metric we used was the number of protein-coding genes separating the genes in each pair. To calculate this distance, we first extracted the genomic positions of all human genes from the human gene-feature table downloaded from the NCBI repository (*78*). Next, we used a custom Perl script to calculate how many protein-coding genes are found between genes in each pair analyzed, and grouped the analyzed gene pairs into the categories described in the Results section.

Additionally, we analyzed all pairs of genes found on the same chromosome together. This group was labeled ‘same chromosome’ gene pairs. The analysis of all ‘same chromosome’ gene pairs together was carried out to see if there is any difference between the fusion-related gene pairs and the controls, before performing analyses of the same-chromosome sub-groups.

In cases of same-chromosome genes where the start position of the downstream gene was upstream to the end position of the upstream gene, the genes were labeled ‘overlapping’. If the whole interval of one of the genes, from start to end, was included in the interval of the other gene, the genes were labeled ‘included’. Gene pairs in the ‘included’ or ‘overlapping’ categories, as well as gene pairs with no protein-coding genes between the pair members, were considered ‘neighbors’ (SC_0). Furthermore, when counting the number of protein-coding genes separating pair members in an analyzed pair, ‘included’ genes were not taken into account, since they do not actually contribute to the distance between the two focal genes more than the larger genes in which they are included.

If one of the genes in a pair was on a non-localized/unplaced scaffold or on the alternate loci assembly, the distance between these two genes was marked as ‘unclear’. These gene-pairs were excluded from further analyses.

### Creating lists of randomized control pairs

To test whether fusion-related gene pairs interact more strongly than random gene pairs, we created two types of control gene-pair lists: genomic control and STRING control. The genomic control was used to test whether the members of fusion-related gene pairs identified using the STRING database tended to interact more strongly than the members of random pairs of genes in the human genome. To prepare this control list, for each pair of fusion-related genes from STRING, we drew a random gene from the human genome using the ‘sample’ function in R (*79*) and then paired this random gene with a gene downstream to it found at the same distance from it as the distance between the members of the focal STRING fusion-related gene-pair (distance was measured by the number of protein-coding genes separating the genes in the pair). If a random gene could not be assigned a partner to form a pair that was distance-matched to the focal fusion-related pair, for example because it was found at the end of the chromosome, this random gene was discarded and another gene was randomly chosen from the genome. Only gene pairs not found in the STRING database were used to create the lists of genomic control pairs, though STRING pairs constitute a minor fraction of all possible random genomic pairs.

The STRING control was used to test whether the members of fusion-related gene pairs identified using the STRING database tended to interact more strongly with each other than members of fusion-unrelated gene pairs from the same database. To prepare this STRING control list, all fusion-unrelated gene pairs in the STRING database (*28*) were grouped by the distance between the members of each pair (as measured by the number of protein-coding genes separating the members). Next, for each fusion-related gene pair we chose distance-matched random STRING gene pairs using the ‘sample’ function in R (*79*).

The process of choosing random gene pairs for both the genomic and STRING controls was repeated ten times per each fusion-related pair, thus producing control lists (both genomic and STRING) 10x larger than the list of fusion-related pairs, providing sufficient statistical power. In cases where the final number of distance-matched random gene pairs was smaller than 10x, for any measured distance, we used all available pairs for the analysis.

### Co-expression analysis

To analyze the co-expression of fusion-related gene pairs compared to random pairs we used the human gene co-expression data downloaded from the COXPRESdb portal (*29, 80*). First, we used a custom Perl script to extract the co-expression score for each pair of fusion-related and control genes. Next, we used the one-sided Mann-Whitney-Wilcoxon test, as implemented by the ‘wilcox.test’ function in R (*79*), to test whether fusion-related gene pairs are more highly co-expressed than random pairs. Note that here, a lower score represents a stronger degree of co-expression.

The COXPRESdb portal has four different databases of human gene co-expression (*80*). The difference between them is in the co-expression platform used to obtain the data and in the methods used to compute the co-expression scores from the raw data (*29*). In our analysis we used all four databases, with each database being analyzed separately, to examine whether the results obtained are consistent across the databases used.

To avoid generating control gene pairs for which information in the database is lacking, only genes for which information is present were used to generate the control lists.

### Co-localization analysis

To compare the co-localization of fusion-related gene pairs to random pairs, we used a database of protein sub-cellular localization predicted by WoLF PSORT, downloaded from the CoexpressDB portal (*29, 30*).

To decide whether the members of a gene pair are co-localized, we implemented the following method of semantic similarity comparison: If any of the protein products of one gene, and any of the protein products of the other gene, were associated with the same cellular compartment term, the gene pair was marked as co-localized. To test whether the proportion of gene pairs whose products co-localize is higher among fusion-related pairs than control pairs we used a one-tailed Fisher exact test, as implemented by the ‘fisher.test’ function in R (*79*).

As for the co-expression analysis, to avoid generating control gene pairs for which information in the database is lacking, only genes for which information is present were used to generate the control lists.

### Topologically associating domain (TAD) presence analysis

The data of TAD coordinates in the human genome downloaded from the 3D Genome Browser portal (*31*) consisted of several independent lists containing TAD coordinates resulting from a number of different experiments and studies. The lists were created by Feng Yue et al. using an in-house pipeline (*31*).

To study the same-TAD presence of the analyzed gene pairs, first we identified the boundaries of the genes in each pair using data from the ‘gene feature table’ of *H. sapiens* downloaded from the NCBI repository (*78*). Next, we determined for each gene pair the number of individual TAD coordinate lists within the 3D Genome Browser database in which both genes in the pair were found in the same TAD. A gene was considered to be present in a TAD if the entirety of it (from start to end) was included in that TAD. Next, we used the one-sided Mann-Whitney-Wilcoxon test, as implemented by the ‘wilcox.test’ function in R (*79*), to examine whether fusion-related gene pairs are found in the same TAD across a larger number of TAD lists than control pairs.

### Analysis of GO term semantic relatedness

To obtain the GO terms associated with the analyzed genes, we downloaded the ‘Gene2GO’ list, which associates GO terms with genes, from the NCBI repository. Next, we used a custom Perl script to extract GO terms for the analyzed genes from this database. Then, we used GOGO, a program for measuring semantic similarity of GO terms (*32*), to obtain the similarity score of GO terms for genes in each analyzed pair. Finally, we tested whether those similarity scores were larger in the fusion-related than control gene pairs using a one-sided Mann-Whitney-Wilcoxon test, as implemented by the ‘wilcox.test’ function in R (*79*)

The analysis was conducted separately for each of the three main categories of GO terms: Biological Process (BP), Molecular Function (MF) and Cellular Component (CC). We analyzed these categories separately because not all studied genes had associated GO terms for all three categories. Accordingly, the control lists for these analyses were also created separately for each category.

### Analysis of presence in the database of cancer-related fusions

To analyze whether human gene pairs fused in other species are also likely to undergo fusion in various human cancers, we downloaded data of gene fusions in cancers from the Cosmic (*33*) and FusionGDB portals (*34, 35*). The Cosmic database is small but highly accurate since its contents are manually curated by a large panel of experts (*33*). FusionGDB provides a large list of gene fusions in cancer by integrating data from three sources (*34, 35*): *a*) the database of chimeric transcripts and RNA-seq data (ChiTaRS 3.1); *b*) an integrative resource for cancer-associated transcript fusions (TumorFusions) and *c*) the Cancer Genome Atlas (TCGA) fusions by Gao et. al. For the purpose of the analysis, data from the two sources were combined into a single list of gene pairs that have been observed to undergo fusion in cancer (whether DNA fusion or fusion of their transcripts).

We used a custom Perl script to calculate the number of gene pairs that overlap between the lists of evolutionary-fusion–related pairs from STRING and gene pairs involved in cancer fusions, as well as between the control gene pairs and the latter. Next, we used a one-sided Fisher exact test, as implemented by the ‘fisher.test’ function in R (*79*) to determine whether the proportion of overlapping pairs was significantly higher among evolutionary-fusion–related than among control gene pairs.

## Notes

### Competing Interest Statement

The authors have declared no competing interest.

## References

1. Virgen CA, Kratovac Z, Bieniasz PD, Hatziioannou T (2008) Independent genesis of chimeric TRIM5-cyclophilin proteins in two primate species. P Natl Acad Sci USA 105:3563–3568.

2. Haendler B, Hofer E (1990) Characterization of the human cyclophilin gene and of related processed pseudogenes. Eur J Biochem 190(3):477–482.

3. Kaessmann H, Vinckenbosch N, Long M (2009) RNA-based gene duplication: mechanistic and evolutionary insights. Nat Rev Genet 10:19–31.

4. Nisole S, Lynch C, Stoye JP, Yap MW (2004) A Trim5-cyclophilin A fusion protein found in owl monkey kidney cells can restrict HIV-1. P Natl Acad Sci USA 101:13324–13328.

5. Sayah DM, Sokolskaja E, Berthoux L, Luban J (2004) Cyclophilin A retrotransposition into TRIM5 explains owl monkey resistance to HIV-1. Nature 430:569–573.

6. Liao CH, Kuang YQ, Liu HL, Zheng YT, Su B (2007) A novel fusion gene, TRIM5-Cyclophilin A in the pig-tailed macaque determines its susceptibility to HIV-1 infection. Aids 21(Suppl 8):S19–S26.

7. Brennan G, Kozyrev Y, Hu SL (2008) TRIMCyp expression in old world primates *Macaca nemestrina* and *Macaca fascicularis*. P Natl Acad Sci USA 105:3569–3574.

8. Wilson SJ, et al. (2008) Independent evolution of an antiviral TRIMCyp in rhesus macaques. P Natl Acad Sci USA 105:3557–3562.

9. Newman RM, et al. (2008) Evolution of a TRIM5-CypA splice isoform in old world monkeys. PLoS Pathog 4:e1000003.

10. Zhang F, Hatziioannou T, Perez-Caballero D, Derse D, Bieniasz PD (2006) Antiretroviral potential of human tripartite motif-5 and related proteins. Virology 353(2):396–409.

11. Yap MW, Dodding MP, Stoye JP (2006) Trim-cyclophilin A fusion proteins can restrict human immunodeficiency virus type 1 infection at two distinct phases in the viral life cycle. J Virol 80(8):4061–4067.

12. Yap MW, Mortuza GB, Taylor IA, Stoye JP (2007) The design of artificial retroviral restriction factors. Virology 365(2):302–314.

13. Livnat A, Papadimitriou C (2016) Evolution and learning: used together, fused together. A response to Watson and Szathmáry. Trends in Ecology & Evolution 31(12):894–896.

14. Livnat A (2017) Simplification, innateness, and the absorption of meaning from context: how novelty arises from gradual network evolution. Evolutionary Biology 44(2):145–189.

15. Jackson DA, Hassan AB, Errington RJ, Cook PR (1993) Visualization of focal sites of transcription within human nuclei. The EMBO Journal 12(3):1059.

16. Edelman LB, Fraser P (2012) Transcription factories: genetic programming in three dimensions. Current Opinion in Genetics & Development 22(2):110–114.

17. Papantonis A, Cook PR (2013) Transcription factories: genome organization and gene regulation. Chemical Reviews 113(11):8683–8705.

18. Marcotte EM, et al. (1999) Detecting protein function and protein-protein interactions from genome sequences. Science 285(5428):751–753.

19. Enright AJ, Iliopoulos I, Kyrpides NC, Ouzounis CA (1999) Protein interaction maps for complete genomes based on gene fusion events. Nature 402(6757):86–90.

20. Stone E, Schwartz R (1990) Intron-dependent evolution of progenotic enzymes in Intervening Sequences in Evolution and Development, eds. Stone E, Schwartz R. (Oxford University Press, New York), pp. 63–91.

21. West-Eberhard MJ (2003) Developmental Plasticity and Evolution. (Oxford University Press).

22. Doolittle RF (1999) Do you dig my groove? Nature Genetics 23(1):6–8.

23. Gingeras TR (2009) Implications of chimaeric non-co-linear transcripts. Nature 461(7261):206.

24. Carvalho CM, Zhang F, Lupski JR (2010) Genomic disorders: A window into human gene and genome evolution. Proc Natl Acad Sci USA 107(suppl 1):1765–1771.

25. Livnat A (2013) Interaction-based evolution: how natural selection and nonrandom mutation work together. Biology Direct 8(1):24.

26. Li H, Wang J, Mor G, Sklar J (2008) A neoplastic gene fusion mimics trans-splicing of RNAs in normal human cells. Science 321(5894):1357–1361.

27. Osborne CS (2014) Molecular pathways: transcription factories and chromosomal translocations. Clinical Cancer Research 20(2):296–300.

28. Szklarczyk D, et al. (2019) STRING v11: protein-protein association networks with increased coverage, supporting functional discovery in genome-wide experimental datasets. Nucleic Acids Res 47(D1):D607–D613.

29. Obayashi T, Kagaya Y, Aoki Y, Tadaka S, Kinoshita K (2018) COXPRESdb v7: a gene coexpression database for 11 animal species supported by 23 coexpression platforms for technical evaluation and evolutionary inference. Nucleic Acids Research 47(D1):D55–D62.

30. Horton P, et al. (2007) Wolf psort: protein localization predictor. Nucleic acids research 35(Web Server issue):W585—7.

31. Wang Y, et al. (2018) The 3D Genome Browser: a web-based browser for visualizing 3D genome organization and long-range chromatin interactions. Genome biology 19(1):1–12.

32. Zhao C, Wang Z (2018) GOGO: An improved algorithm to measure the semantic similarity between gene ontology terms. Scientific reports 8(1):1–10.

33. Tate JG, et al. (2018) COSMIC: the Catalogue Of Somatic Mutations In Cancer. Nucleic Acids Research 47(D1):D941–D947.

34. Kim P, Zhou X (2019) FusionGDB: fusion gene annotation DataBase. Nucleic Acids Res 47(D1):D994–D1004.

35. Kim P, Zhou X (2018) “FusionGDB: Fusion Gene annotation DataBase” https://ccsm.uth.edu/fusiongdb Accessed 1/24/2019.

36. Kummerfeld SK, Teichmann SA (2005) Relative rates of gene fusion and fission in multi-domain proteins. Trends in Genetics 21(1):25–30.

37. Michalak P (2008) Coexpression, coregulation, and cofunctionality of neighboring genes in eukaryotic genomes. Genomics 91(3):243–248.

38. Koonin EV (2009) Evolution of genome architecture. The International Journal of Biochemistry & Cell Biology 41(2):298–306.

39. Ghanbarian AT, Hurst LD (2015) Neighboring genes show correlated evolution in gene expression. Molecular Biology and Evolution 32(7):1748–1766.

40. Lian S, et al. (2018) Intrachromosomal colocalization strengthens co-expression, co-modification and evolutionary conservation of neighboring genes. BMC Genomics 19(1):1–13.

41. Lynch M (2007) The Origins of Genome Architecture. (Sinauer Associates Sunderland).

42. Gilbert W (1978) Why genes in pieces? Nature 271(5645):501.

43. Fischer SE, Butler MD, Pan Q, Ruvkun G (2008) Trans-splicing in *C. elegans* generates the negative RNAi regulator ERI-6/7. Nature 455(7212):491–496.

44. Graur D, Li WH (2000) Fundamentals of Molecular Evolution, 2nd ed. (Sinauer Associates, Sunderland, MA).

45. Fisher RA (1930) The Genetical Theory of Natural Selection. (The Clarendon Press, Oxford).

46. Kimura M (1968) Evolutionary rate at the molecular level. Nature 217:624–626.

47. King J, Jukes T (1969) Non-Darwinian evolution. Science 164:788–798.

48. Ohno S (1972) So much ’junk’ DNA in our genome in Evolution of Genetic Systems, Brookhaven Symp. Biol. pp. 366–370.

49. King MC, Wilson AC (1975) Evolution at two levels in humans and chimpanzees. Science 188(4184):107–116.

50. Ohta T (2002) Near-neutrality in evolution of genes and gene regulation. Proceedings of the National Academy of Sciences 99(25):16134–16137.

51. Melamed D, et al. (2021) *De novo* mutation rates at the single-mutation resolution in a human HBB gene-region associated with adaptation and genetic disease. bioRxiv.

52. Ord TJ, Summers TC (2015) Repeated evolution and the impact of evolutionary history on adaptation. BMC Evolutionary Biology 15(1):1–12.

53. Conte GL, Arnegard ME, Peichel CL, Schluter D (2012) The probability of genetic parallelism and convergence in natural populations. Proceedings of the Royal Society B: Biological Sciences 279(1749):5039–5047.

54. Blount ZD, Lenski RE, Losos JB (2018) Contingency and determinism in evolution: Re-playing life’s tape. Science 362(6415).

55. Xie KT, et al. (2019) DNA fragility in the parallel evolution of pelvic reduction in stickleback fish. Science 363(6422):81–84.

56. Kratochwil CF, Liang Y, Urban S, Torres-Dowdall J, Meyer A (2019) Evolutionary dynamics of structural variation at a key locus for color pattern diversification in cichlid fishes. Genome Biol Evol 11(12):3452–3465.

57. Kratochwil CF, Meyer A (2019) Fragile DNA contributes to repeated evolution. Genome Biol 20(1):39.

58. Lind PA (2019) Repeatability and predictability in experimental evolution in Evolution, Origin of Life, Concepts and Methods, ed. Pontarotti P. (Springer), pp. 57–83.

59. Flint J, Harding RM, Boyce AJ, Clegg JB (1998) The population genetics of the haemoglobinopathies. Baillière’s Clin Haem 11:1–51.

60. Snel B, Bork P, Huynen M (2000) Genome evolution: gene fusion versus gene fission. Trends in Genetics 16(1):9–11.

61. Nakamura Y, Itoh T, Martin W (2007) Rate and polarity of gene fusion and fission in *Oryza sativa* and *Arabidopsis thaliana*. Molecular Biology and Evolution 24(1):110–121.

62. Leonard G, Richards TA (2012) Genome-scale comparative analysis of gene fusions, gene fissions, and the fungal tree of life. Proceedings of the National Academy of Sciences 109(52):21402–21407.

63. Lupski JR (1998) Genomic disorders: structural features of the genome can lead to DNA rearrangements and human disease traits. Trends Genet 14:417–422.

64. Gu W, Zhang F, Lupski JR (2008) Mechanisms for human genomic rearrangements. Patho-Genetics 1:4.

65. Woodward KJ, et al. (2005) Heterogeneous duplications in patients with pelizaeus-merzbacher disease suggest a mechanism of coupled homologous and nonhomologous recombination. Am J Hum Genet 77:966–987.

66. Lee JA, et al. (2006) Role of genomic architecture in *PLP1* duplication causing pelizaeus-merzbacher disease. Hum Mol Genet 15:2250–2265.

67. Hastings PJ, Ira G, Lupski JR (2009) A microhomology-mediated break-induced replication model for the origin of human copy number variation. PLoS Genet 5:e1000327.

68. Korbel JO, et al. (2007) Paired-end mapping reveals extensive structural variation in the human genome. Science 318:420–426.

69. Lupski JR (2006) Genome structural variation and sporadic disease traits. Nature Genet 38:974–976.

70. Williams GC (1966) Adaptation and Natural Selection. (Princeton University Press).

71. Dawkins R (1976) The Selfish Gene. (Oxford University Press).

72. Ohno S (1970) Evolution by Gene Duplication. (Springer-Verlag, Heidelberg).

73. Jacob F (1977) Evolution and tinkering. Science 196:1161–1166.

74. Hebb D (1949) The Organization of Behavior. (Wiley & Sons, New York).

75. Lindley R (1966) Recording as a function of chunking and meaningfulness. Psychonomic Science 6:393–394.

76. Löwel S, Singer W (1992) Selection of intrinsic horizontal connections in the visual cortex by correlated neuronal activity. Science 255(5041):209–212.

77. Tulving E, Craik FI (2005) The Oxford Handbook of Memory. (Oxford University Press).

78. O’Leary NA, Wright MW, Brister JR, , et al. (2016) Reference sequence (RefSeq) database at NCBI: current status, taxonomic expansion, and functional annotation. Nucleic Acids Res 44(D1):D733–745.

79. R Core Team (2019) R: A Language and Environment for Statistical Computing (R Foundation for Statistical Computing, Vienna, Austria).

80. Obayashi T, Kagaya Y, Aoki Y, Tadaka S, Kinoshita K (2019) “COXPRESdb” https://coxpresdb.jp Accessed 03/18/2019.

